# Human Dental Pulp Stem Cells Grown in Neurogenic Media Differentiate into Endothelial Cells and Promote Neovasculogenesis in the Mouse Brain

**DOI:** 10.1101/467035

**Authors:** J. Luzuriaga, O. Pastor-Alonso, J.M. Encinas, F. Unda, G. Ibarretxe, JR. Pineda

## Abstract

Dental Pulp Stem Cells (DPSCs) have a demonstrated capacity to acquire neuronal-like phenotypes, suggesting their use in brain cell therapies. In the present work, we wanted to address the phenotypic fate of adult DPSCs cultured in Neurocult media (Stem Cell Technologies), a cell culture medium without serum routinely used for the expansion of adult neural stem cells (NSCs). Our results showed for the first time, that non-genetically modified adult DPSCs cultured with Neurocult generated neurosphere-like dentospheres expressing the NSC markers Nestin and GFAP, but also the vascular endothelial cell marker CD31. One month post-intracranial graft into athymic nude mice, human CD31+ or Nestin+ DPSC-derived cells were found tightly associated with brain blood vessels increasing their laminin staining. These results suggest that DPSCs integrated and contributed to an increased generation of neovasculature within brain tissue and that Neurocult medium constituted a fast and efficient way to obtain endothelial cells from human DPSCs.

## INTRODUCTION

The dental pulp is home to a surprisingly active population of neural crest stem cells, which are responsible of renewing adult differentiated cells, such as odontoblastic cells and Schwann cells, within the pulp tissue^1, 2^. These cells are referred to as Dental Pulp Stem Cells (DPSCs) when extracted from permanent teeth, and as Stem cells from Human Exfoliated Deciduous Teeth (SHEDs) when extracted from first generation teeth during childhood^3-5^. Periodontal tissues also constitute a very rich source of stem cells with related characteristics^6-8^. All the aforementioned cell populations share stemness properties^3, 4^ and they are able to differentiate into several cell types, preferentially to those of mesenchymal lineages, like osteoblasts, chondroblasts and adipocytes^3, 9^. Differentiation of SHEDs and DPSCs towards neural lineages has also been extensively reported in the literature in the presence^10^ or absence of scaffolds^11-13^, especially when cells were cultured with DMEM/F-12 or Neurobasal medium supplemented with B27 and EGF/FGF growth factors^14-17^, although, a large debate remains as to whether those cells are genuinely neuronal or merely neuronal marker-expressing cells (neuron-like), or whether they can effectively integrate into host brain neuronal network after their graft *in vivo*.

Interestingly, DPSCs also have the ability to co-differentiate synergically to osteoblast and endothelial cell phenotypes when cultured with Fetal Bovine Serum (FBS)-containing media^18, 19^. The possibility to use these cells as a source of young neovasculature has not been considered, despite the generation of a highly vascularized bone-like tissue after their subcutaneous graft *in vivo*^18, 19^. One of the limitations is the high presence of fetal serum (10 - 20 % FBS + α-MEM)^16, 18, 20^. Fetal serum permits cell survival and rapid cell expansion for the subsequent application of differentiation protocols. However, its presence also favors differentiation toward osteoblastic/odontoblastic lineages^21, 22^. Moreover, serum incorporation during stem cell culture procedure might also cause allergies and immune reactions of the transplanted cells^23^.

In this study, we explored the possibility to obtain endothelial cells ready to use for *in vivo* grafts starting from genetically unmodified human DPSCs extracted and cultured directly with completely serum-free medium. For this purpose we chose Neural Stem Cell media supplemented with B27, heparin, and growth factors. This culture medium is an alternative method for DPSC expansion compatible with the *in vitro* differentiation to neuronal, astroglial (neuroectoderm) cells, but also is permissive for the production of endothelial cells (mesoderm), without a need of scaffolds or serum-addition. Moreover, the *in vivo* differentiation/fate of DPSC grafts when transplanted into the brain of immunocompromised nude mice demonstrates a full differentiation towards endothelial lineage without osteoblast/cartilage production. The clinical relevance is major, because (i) we obtained an enrichment of endothelial cell production starting from DPSC free floating dentospheres and (ii) we do not use at any step fetal serum, known to be highly allergenic and responsible for the rejection of cell transplants in humans^23^.

## RESULTS

### Characterization of DPSC growth dynamics and neural stem marker expression using Neurocult proliferation media

Previous research had shown that the dental pulp tissue of adult teeth contained neural-crest derived DPSCs, which expressed neural stem cell markers such as Nestin and GFAP^4, 11^. Moreover, these were fully viable after one month post-graft and capable of *in vitro* differentiating into mesenchymal cell lineages and generating a dentin pulp-like tissue complex, after *in vivo* subcutaneous transplantation^3^. Our first aim was to evaluate whether Neurocult, a serum-devoid cell culture medium, widely used to grow adult neural stem cells and progenitors, would be also permissive for the growth of DPSCs. For this purpose, we compared DPSC cultures grown with the standard media DMEM/FBS and Neurocult proliferation media. Interestingly DPSCs cultured with Neurocult proliferation media maintained the expression of the brain-derived neurothropic factor (*BDNF*. Supplementary Fig. S1), a neurotrophin involved in neurogenesis and neuron survival^24, 25^. As Neurocult proliferation medium is specially designed to generate neurospheres from dissected and isolated brain neurogenic niches, we used control cultures of neural stem cells from mouse hippocampi, by adapting the protocol previously described^26^.

Both DPSC-derived dentospheres and NSC-derived neurospheres were grown in Neurocult proliferation media. Once disaggregated from spheres, both NSCs and DPSCs were induced to attach to the platting surface using laminin-coated coverslips for 45 minutes, and they were immunostained against the intermediate filament Nestin, a marker expressed by neural stem cells both *in vitro* and *in vivo*^27, 28^. Both DPSCs grown with DMEM + 10% FBS or Neurocult Proliferation media cells were positive for Nestin, as well as NSC (Figure 1AB).

**Figure 1:**
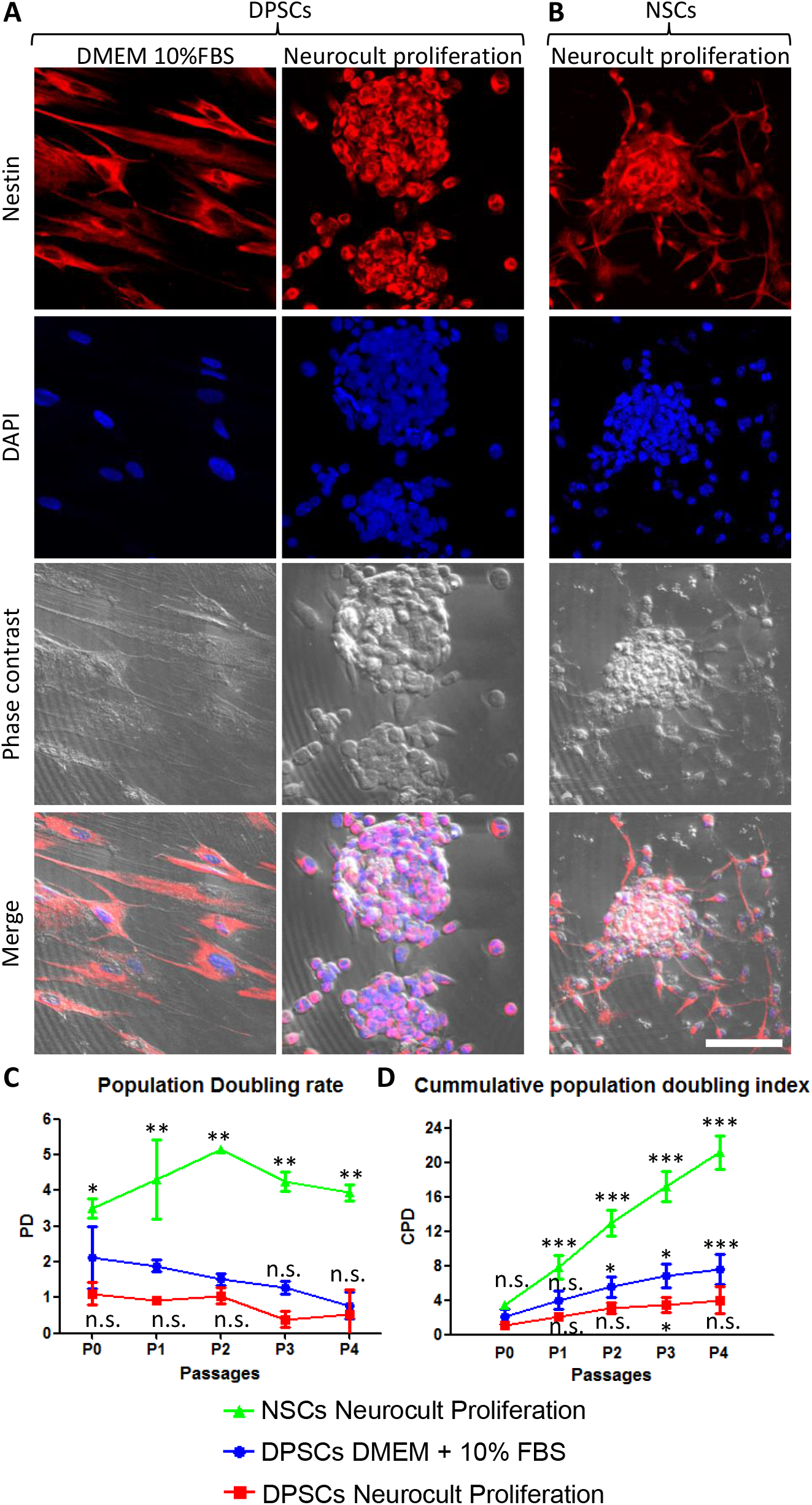
Human DPSCs grown in Neurocult proliferation culture media generate Dentospheres similar to Neurospheres from murine Neural Stem Cells (NSCs). **A)** Human DPSCs seeded in standard culture DMEM + 10% FBS acquire a flat morphology and adhere to the flask surface. However, when cultured with Neurocult proliferation media **(B)** DPSCs start to grow forming free-floating dentospheres. **(C)** Control neurospheres of murine NSCs. All cells are positive for the Neural Stem/progenitor marker Nestin. **(D)** Population Doubling rates for human DPSCs grown either with DMEM + 10% FBS or Neurocult proliferation compared with murine NSCs. **E**) Cumulative population doubling (CPD) of DPSCs seeded in standard culture DMEM+10%FBS or Neurocult proliferation media and NSCs cultured in Neurocult proliferation media. All different cell culture conditions were assessed in parallel (n=3 for each) for a total of 4 passages and non-parametric Kruskal-Wallis with post-hoc test was used (Mean±SD *p<0.05, **p<0.01 and ***p<0.001). Scale bar 75 µm.

We evaluated the growth kinetics of DPSCs cultured either using DMEM + 10% FBS or Neurocult proliferation, and NSCs grown with Neurocult proliferation media. Human DPSCs showed less proliferative index than murine NSC (Figures 1C-D). The population doubling rate (PD) showed no statistical significant differences between Neurocult proliferation media respect to DMEM + 10% FBS but faster growth of NSCs (p=0.0556 and p=0.0037 respectively. One-tailed Kruskal Wallis test; Figure 1C). However, the analysis of cumulative population doubling (CPD) showed that both DPSCs cultures had a parallel trend with a steadily rising growth kinetics, after consecutive passages without significant differences until P3, where differences began to emerge between both conditions (p=0.0273, one-tailed Kruskal Wallis test; Figure 1D). It is noteworthy that without coating, DPSCs cultured using DMEM + 10% FBS grew attached to the surface. Meanwhile DPSCs in Neurocult proliferation media had a progressively slower growth, and generated free-floating dentospheres during all the successive passages, as normally NSCs do.

Triple immunofluorescence analysis was performed against GFAP, Nestin and S100β at 3 days post seeding for high proliferating NSCs, and at 7 days for the more slow growing human DPSCs in Neurocult proliferation media. Both cultures showed comparable proportions of label-positive cells in DPSCs and NSCs, for each of the three tested markers (Figure 2). Nestin-GFAP coexpression is a long-known and widely reported marker profile characteristic of NSCs^26, 29^, and this also has been found in DPSCs cultures using neuronal inductive media^11^. The mature astrocytic marker S100β, whose expression in GFAP-expressing cells coincides with the loss of their NSCs potential^30^, could also be detected in a minority of cells, both in DPSC and NSC cultures. We determined that all human DPSCs cells were Nestin positive, in agreement with the results published by Chang et al.^11^ with a proportion of 69 ± 34 % GFAP and 24 ± 17 % S100β positive cells (Figure 2A-A’). 96±18 % murine NSCs grown with Neurocult proliferation media were Nestin-positive with a proportion of 52±12 % GFAP and 35±10 % S100β positive cells (Figure 2B-B’). Thus, DPSCs and NSCs grown in Neurocult proliferation media contained similar proportions of cells expressing neural stem cell markers, and putatively astrocyte-committed S100β positive cells.

**Figure 2:**
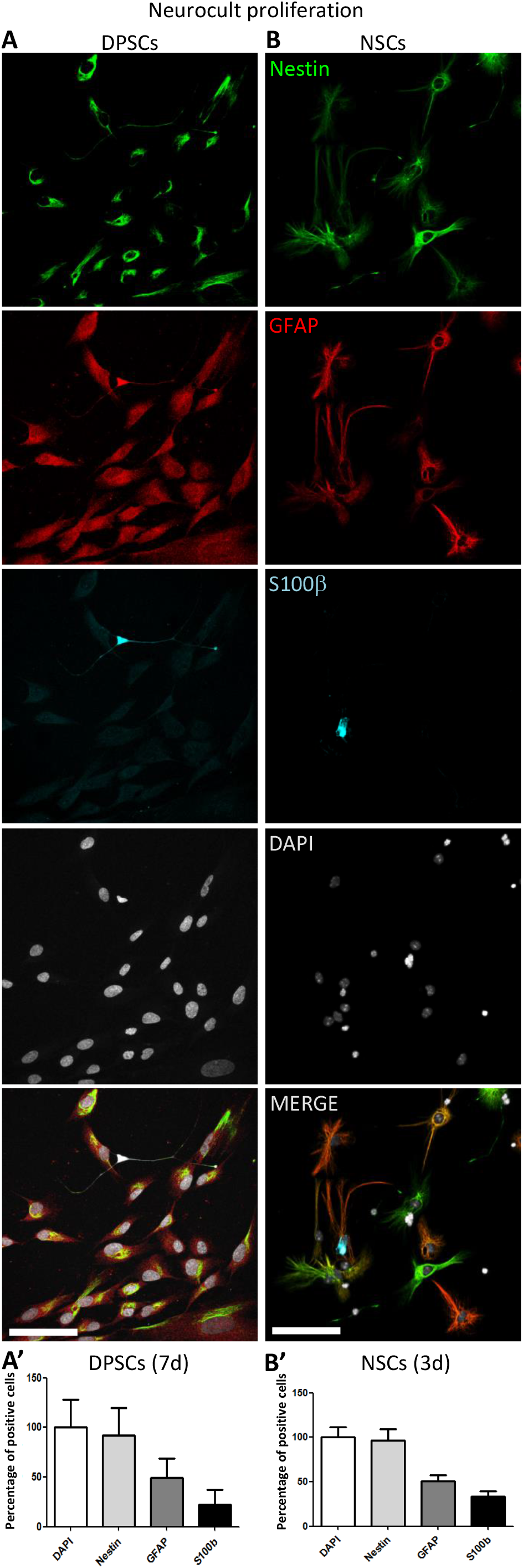
Human DPSCs grown in Neurocult proliferation media share neural stem/progenitor markers with murine NSCs. **A-A’)** Human DPSCs allowed to grow for 7 days in laminin-coated wells in the presence of Neurocult proliferation media are Nestin positive and also express astroglial markers GFAP and S100ß (n=230). **B-B’)** Murine NSCs grown for 3 days (to avoid confluency) in the same culture conditions also express these astroglial markers in similar proportion (n=242). Mean±SEM of three independent experiments. Scale bar 75 μm.

### DPSCs grown using Neurocult proliferation media increased CD31 expression

SHEDs and DPSCs grown in the presence of serum are able to differentiate into endothelial vasculature^18, 31, 32^, but it was not known whether this capability was dependent or not on the presence of serum. Given that DPSCs grown in serum-free Neurocult proliferation media expressed the stem cell marker Nestin, which has also been reported as a marker of proliferative endothelial cell progenitors ^33^, we wondered if serum-free Neurocult proliferaton would also be permissive for the generation of endothelial cells by detecting expression of *CD31* and *VEGF*, both markers of endothelial cells^34, 35^. Q-PCR of *CD31* mRNA expression in DPSCs cultured with Neurocult proliferation showed an increase up to several orders of magnitude (249,57±172,2 fold-increase of expression) with respect to cultures with DMEM 10% FBS (p=0.0286; Mann Whitney test, Figure 3A). To confirm this result, in a next step we decided to perform immunofluorescence analysis for the endothelial markers CD31 and VEGF on laminin-coated slides in both DPSC grown in DMEM + 10% FBS and Neurocult and count the proportion of positive cells. Surprisingly, a generalized nuclear staining for VEGF was found in DPSCs for both culture conditions (Figure 3B), which was confirmed by PCR detection (Figure 3C). Interestingly, although murine NSCs derived from neurogenic niches also expressed VEGF, they were completely negative for CD31 staining (Figure 3B right panel) and did not integrate into murine vasculature after intracerebral graft in consanguine mice, although they showed some projecting contacts to the brain vasculature, as previously described^26^ (Supplementary Fig. S2). However, it is known that laminin could induce a morphologic differentiation of DPSC to endothelial cells^36^. In order to exclude this possibility, we cultured DPSCs without laminin as floating dentospheres and thereafter we run a flow cytometry analysis by labeling the cells either with CD31-FITC or the IgG control isotype. We detected an increase of up to 26% of CD31-FITC positive cell population in DPSCs when these cells were grown with Neurocult proliferation (Figure 3E *right*). In contrast, the flow cytometry analysis only gave a residual value of 0.3 % of CD31-positive DPSCs, when these cells were grown in DMEM + 10% FBS (Figure 3E *left*). In conclusion, our results demonstrated that serum-free Neurocult proliferation medium is able to increase the expression of the CD31 endothelial cell marker in DPSC cultures.

**Figure 3:**
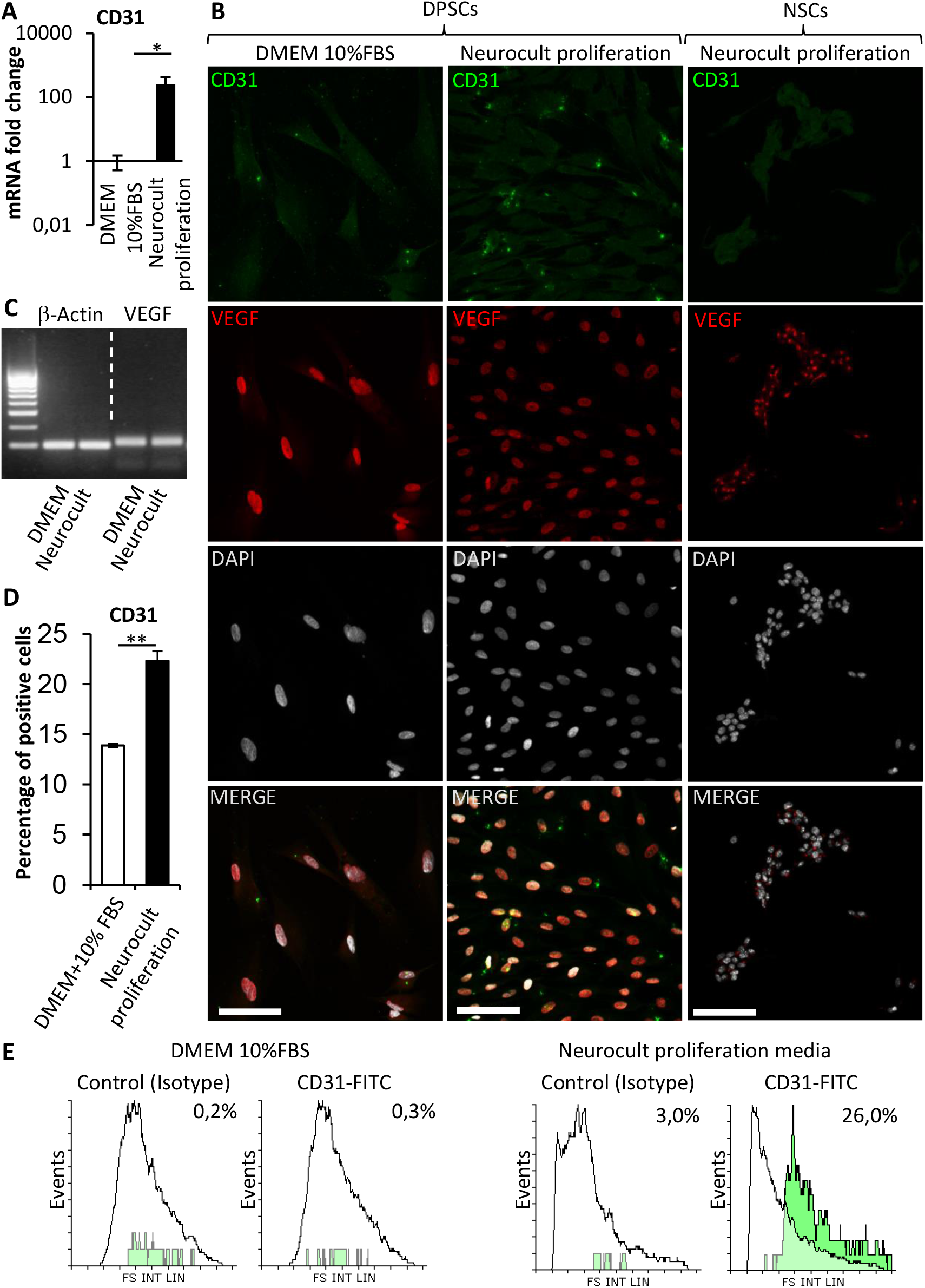
The endothelial CD31 marker is absent from NSCs whereas it can be enriched upon the type of culture media in DPSCs. **A)** mRNA fold change of the endothelial marker CD31 on DPSC cultures upon the type of media: DMEM/FBS vs. Neurocult proliferation. Axis represented as logarithmic scale of CD31 expression from DPSC cultures of 4 different patients (Mean±SEM *p=0.05, one-tailed Mann Whitney test). **B)** Immunofluorescence of DPSCs cultured with (*left column*) DMEM + 10% FBS or (*middle column)* Neurocult proliferation media for CD31 and nuclear VEGF. NSC (*right column*) cultured with Neurocult proliferation. Scale bars 75 μm. **C)** PCR detection for VEGF confirming its expression in DPSCs. Quantification of the percentage of positive cells for CD31 staining by **D)** immunofluorescence (Mean±SEM n=180 for DMEM 10% FBS and n=1564 for Neurocult proliferation, **p=0.01, one-tailed Mann Whitney test), and **E)** Flow cytometry analysis for CD31 expression in DPSC cultures (n=10.000 events for each condition. *p=0.05, one-tailed Mann Whitney test).

### DPSCs grown in Neurocult proliferation media expressed the VEGFR2 receptor and showed an increased ERK pathway activity

It has been previously reported that ERK signaling and VEGFR expression are required for endothelial differentiation of SHED^37^. Because DPSCs grown with Neurocult proliferation media expressed endothelial markers VEGF and CD31, we decided to test the presence of VEGFR2 receptor at mRNA and protein level by Q-PCR and immunofluorescence. DPSCs cultured with Neurocult proliferation showed 17,9±10,2 fold-increase of *VEGFR2* mRNA with respect to cultures with DMEM 10% FBS (p=0.0002; one tailed Mann Whitney test, Figure 4A) and a corresponding increase of VEGFR2 immunostaining (p=0.0125, one tailed Mann Whitney test, Figure 4B-C). Phosphorylation of ERK1/2 protein had been previously reported to be associated to endothelial cell differentiation of SHED^37^ and cell proliferation in a large variety of cell types^38^. In agreement, a specific pERK staining on DMEM 10%FBS DPSC cultures could only be detected on mitotic cells (Figure 4C *left*). In contrast, DPSCs cultured with Neurocult proliferation media clearly showed an increased pERK cytoplasmic staining (Figure 4D *right*). No staining at all was found in controls without primary antibody (Supplementary Fig. 3A). Phospho-ERK and total ERK were quantified by western blot, liver sinusoidal endothelial cells (LSEC) were used as positive control of endothelial cells. Free-floating DPSCs dentospheres cultured with Neurocult proliferation media increased ERK signaling with respect to standard DMEM + 10% FBS conditions (p=0.04 (DMEM + 10% FBS vs Neurocult proliferation) and p=0.0273 (DMEM + 10% FBS vs LSEC); one-tailed Kruskal-Wallis test). In conclusion, DPSCs grown using Neurocult proliferation media increased the activity of the ERK signaling pathway, which had been already described to be essential for SHED to differentiate into endothelial cells.

**Figure 4:**
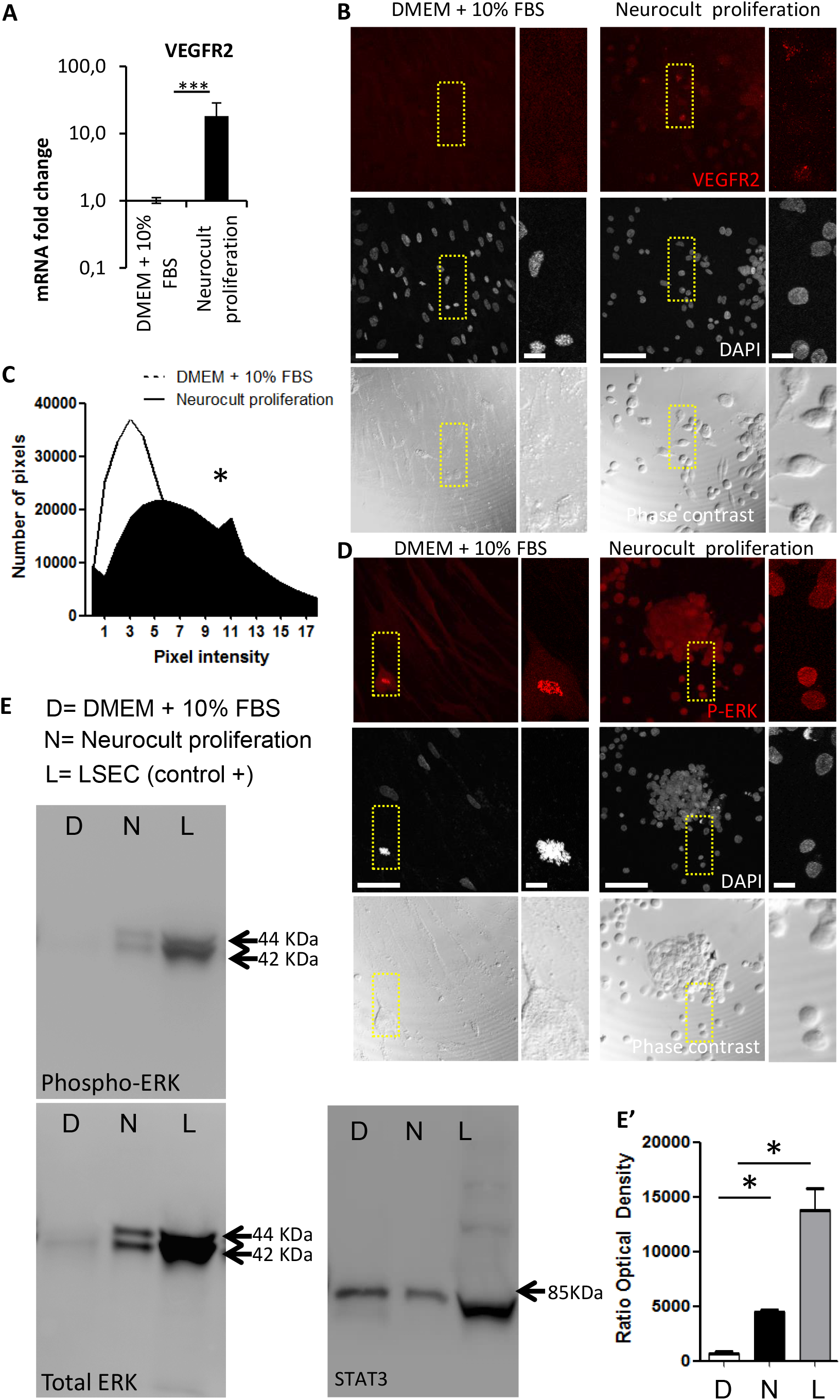
DPSCs grown using Neurocult Proliferation media increase the expression of VEGFR2 and activate downstream ERK signaling. **A)** mRNA fold change of the endothelial marker *VEGFR2* on DPSC cultures depending upon the type of media: DMEM/FBS vs. Neurocult proliferation. Axis represented as logarithmic scale (Mean±SEM of three independent samples, ***p=0.0002, one-tailed, Mann Whitney test). **B)** Immunofluorescence of VEGFR2 receptor staining for cells grown during 7 days with DMEM + 10% FBS (*left*) or Neurocult proliferation (*right*). Scale bar 75 μm, inset 10 μm. **C)** Quantification of distribution of brightest pixel intensity corresponding to VEGFR2 staining (n=139 cells for DMEM + 10% FBS and n=651 cells for Neurocult proliferation. *p=0.0125, one tailed, Mann Whitney test). **D)** Immunofluorescence of phospho-ERK1/2 showed an increase of cytoplasmic staining for cells grown with Neurocult proliferation media, in contrast to the nuclear staining found in mitotic DMEM + 10% FBS DPSCs. Scale bar 75 μm, inset 10 μm. **E)** Western blot of 200.000 cells for each condition showing phospho-ERK1/2 and total ERK protein levels for DPSCs grown either with DMEM + 10% FBS (D) or Neurocult proliferation (N) media and positive control human liver sinusoidal endothelial cells (L, LSEC). STAT3 was used as loading control (full blotting of the gel is shown in Supplementary). **E’)** Quantification of phospho-ERK1/2 western blot in three different samples for each type of culture media (Mean±SEM; *p=0.04 (D vs N) and *p=0.0273 (D vs L); one-tailed Kruskal-Wallis test).

### DPSCs grown in Neurocult differentiation media expressed neuronal and astroglial markers but not endothelial CD31

DPSC cultures using Neurocult proliferation media showed increased levels of VEGFR2, CD31+ and phospho-ERK positive cells. Similarly, a large body of literature suggested that DPSCs were able to differentiate towards neuronal and astroglial lineages in response to the appropriate stimuli^11, 17, 39^. In order to test this hypothesis in our Neurocult media, NSCs and DPSCs were grown in parallel with Neurocult differentiation medium in sphere-disaggregated isolated cells, plated in laminin-coated coverslips (in adherent form) for 7 consecutive days. Neurocult differentiation medium was already known to induce neuronal and astroglial differentiation of murine NSCs^40, 41^. These cells were thus included in the experiment as a positive control for differentiation. To assess neuronal differentiation, we used Doublecortin (DCX) and NeuN markers, both for immature and mature neurons respectively (Figure 5A). On the one hand, when observing the phenotype of differentiated NSCs, we found that 40±8 % of cells were positive for DCX and 36±9 % for nuclear NeuN staining, and differentiated cells showed a characteristic neuron-like dendritic morphology. On the other hand, when neuronal marker expression was assessed in differentiated DPSCs, 39±33 % of cells were positive for DCX, and 34±13 % of them showed positive NeuN staining. However, the morphology of differentiated DPSCs was completely different to NSCs, with long lamellipodia, and neither presence of dendrites nor any other morphological feature that could make them resemble a neuron. Despite a lack of morphological relationship, we did not find any significant difference between the obtained rates of DCX (or NeuN positive cells (p=0.9701 and p=0.8920, respectively; Student’s t test) depending on the type of source cells employed for differentiation: Both DPSCs and NSCs gave rise to neuronal marker-expressing cells in similar proportions (Figure 5B). In addition, the proportions of cells that differentiated towards astroglial lineage were assessed. In this context, Glial Fibrillary Acidic Protein (GFAP) and S100β were used as markers for immature and mature astrocytes, respectively. It was found that 48±15 % of NSCs were positive for GFAP, and 27±7 % for S100β staining. On the other hand, 35±6 % of human DPSCs showed GFAP staining, and only 9±7 % of differentiated DPSCs showed S100β staining, but all cells that did show labeling for S100β also presented a marked bipolar morphology. Again, there were no statistically significant differences in astroglial marker labeling proportions between differentiated DPSCs and NSCs (p=0.0764 Student’s t test, Figure 5C-D). Finally, we decided to check the ratio of endothelial cells. When murine NSC were cultured for seven days in the presence of Neurocult differentiation medium, no expression of the CD31 receptor could be detected but cytoplasmic VEGF staining was still present. In the DPSC counterparts, a dramatic reduction of CD31 labeling was found, with a scarce 5±1 % of CD31-positive DPSCs, when these cells were grown in Neurocult differentiation medium (Figure 5F). However, VEGF expression remained in the nucleus. In conclusion, Neurocult proliferation medium induced human DPSCs, but not murine NSCs, to express endothelial markers. Also, Neurocult differentiation medium induced both DPSCs and NSCs to generate cells which expressed both immature and mature neuronal and astroglial markers in similar proportions, but with very different morphologies, and a near-absence of expression of CD31.

**Figure 5:**
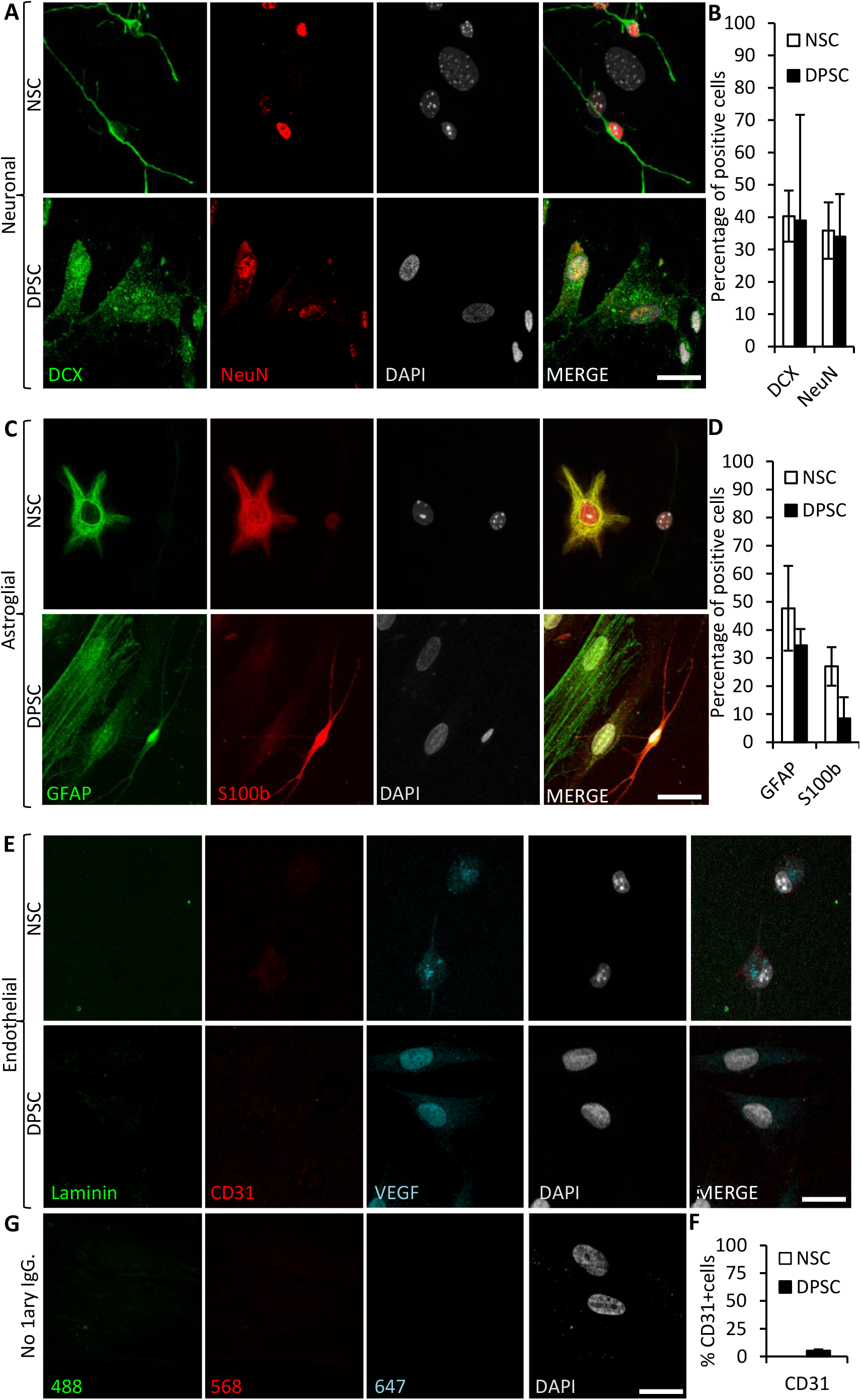
Human DPSCs are able to commit toward differentiation to neuronal-like and glial-like lineages. One week of culture of both human DPSCs and murine (control) NSCs in Neurocult differentiation medium is sufficient to induce them express markers for **(A)** Neuronal lineage differentiation: Doublecortin (DCX) and NeuN staining, for immature and mature neurons respectively, and **C)** Astroglial lineage differentiation: Glial Fibrillary Acidic Protein (GFAP) and S-100β immunostaining, for immature and mature astrocytes, respectively. **B & D)** Graphs showing quantifications (Mean±SEM, n=360 cells) of three independent experiments. **E)** After one week of growth in neural differentiation conditions both NSCs and DPSCs still express VEGF but downregulate CD31. **F)** Quantification of the proportion of CD31 positive cells (n=571). **G)** Control with no 1^ary^ antibodies. Scale bar 20 μm.

### DPSCs grafted *in vivo* migrated and integrated into brain vasculature

In view of the previous *in vitro* results about DPSCs being competent to differentiate to endothelial marker-expressing cells using Neurocult proliferation media, or switch to neuronal and astroglial marker-expressing cells in Neurocult differentiation media, it was important to assess the fate of DPSCs after their *in vivo* graft into the brain. Because no previous data existed about the generation of brain neovasculature from DPSCs, we decided to carry out *in vivo* intracranial cell grafts using DPSCs grown in Neurocult proliferation media. The athymic nude mice model was chosen because it had been previously employed successfully to perform intracranial grafts of human periodontal ligament-derived cells^20^. Ten thousand human DPSC cells were grafted as previously described^26^ varying the stereotaxic coordinates to place cells intrahippocampally. Cell integration in brain tissue was assessed after 30 days as previously described^26^ to better assess the long-term viability of grafted DPSC-derived cells. Human cells were identified using the specific anti-human Nestin antibody as previously described^42^. Grafted cells were located within or very close to blood vessels, which were labeled by Laminin and VEGF staining (Figure 6A). Interestingly, some of the blood vessels appeared to be mostly generated by human DPSC-derived cells, showing a very uniform and homogeneous labeling in coronal sections. Close detail allowed us to determine an increased staining of laminin and VEGF in blood vessels containing human cells, with respect to blood vessels only containing the natural murine vasculature (Figure 6B). To further confirm this result, we performed immunofluorescence against CD31, using an antibody that recognized both human and murine epitopes. This labeling revealed the whole murine brain vasculature, and we found a very high level of colocalization between nestin-positive human DPSC-derived cells, and vascular endothelial CD31-positive cells (Figure 6C). It had been previously reported that Nestin is expressed in both stem cells and proliferative endothelial cells, but not in the mature vasculature^33^. To further validate the presence of human endothelial cells derived from DPSC grafts, we proceeded to label serial sections with a human-specific anti-CD31 antibody. Again, we corroborated the presence of human CD31 positive cells integrated into the host brain vasculature of athymic nude mice at one-month post graft (Figure 6D). Remarkably, with the detection of human cells by specific Nestin or CD31 antibodies, all the vasculature that contained human cells expressed higher amounts of laminin staining. This was quantified by choosing blood vessels of the same caliber, and normalizing the level of intensity of murine vasculature to 100±12 %. Thus, we determined that the presence of human DPSC-derived cells within the blood vessel raised the laminin-labeling intensity to 266±33 % (p<0.0001 Student’s t test, Figure 6E). In summary, we demonstrated that human DPSCs, grown with serum-free Neurocult proliferation medium, were able to survive for one-month and functionally integrate within the brain vasculature of athymic nude mice, expressing markers of endothelial cells.

**Figure 6:**
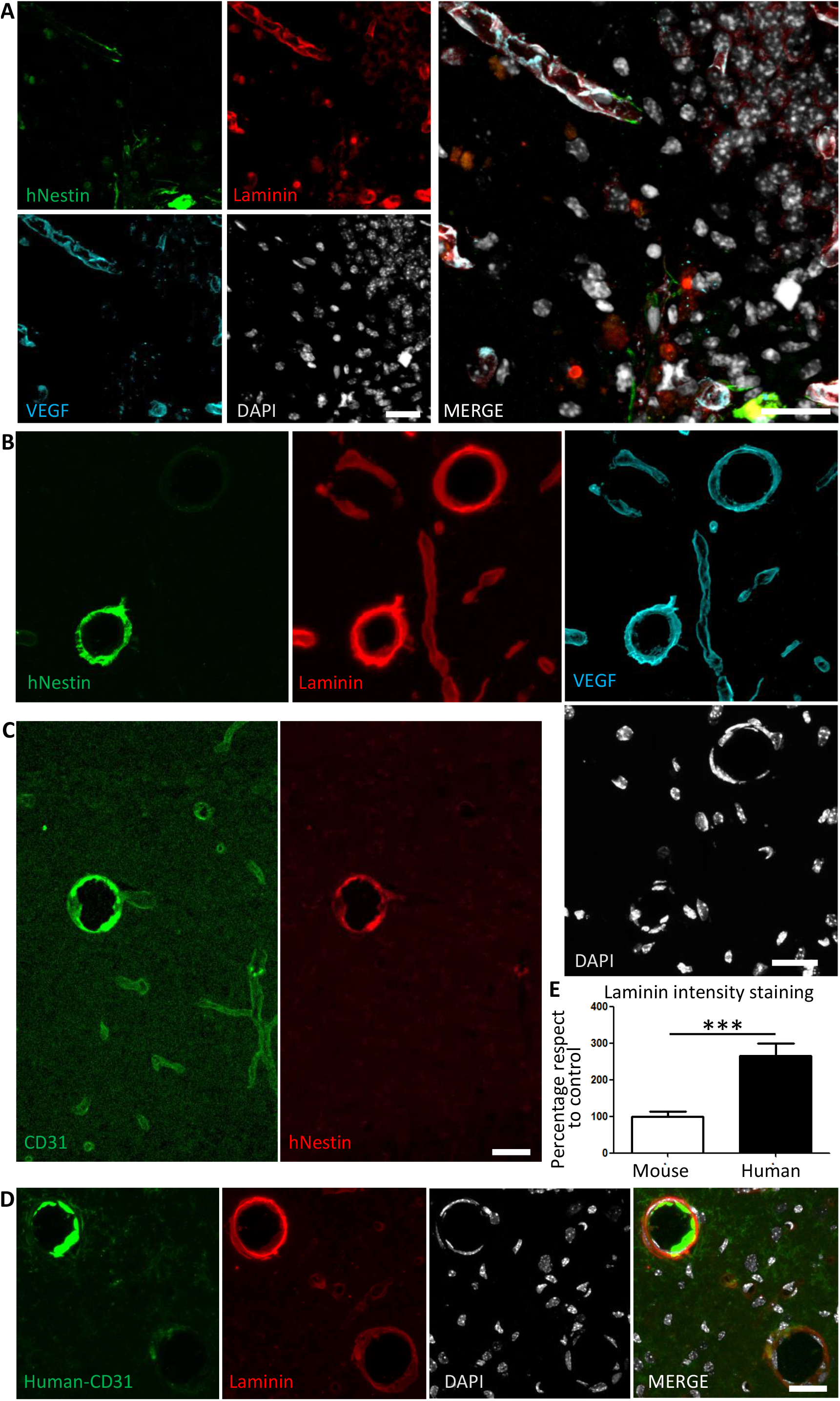
Human DPSCs survive after one-month post graft into the brain of nude mice, expressing CD31 and VEGF and integrating into the host brain vasculature. **A)** One month post-graft into the hippocampus of athymic nude mice, DPSCs are identified by specific human Nestin labeling. Human cells migrate towards Laminin and VEGF-labeled murine vasculature. **B)** Close detail of human Nestin-positive DPSC-derived cells covering brain vasculature showed an increase in the staining for Laminin and VEGF, suggesting *de novo* vasculature formation. **C)** CD31 staining for both murine and human endothelial cells colocalizes with human Nestin-positive DPSC-derived cells in some blood vessels. **D)** A specific antibody for human-CD31 cells recognizes human DPSC-derived endothelial cells in blood vessels, and those vessels precisely correspond to the ones showing an increased laminin staining. **E)** Quantification of laminin staining intensity for murine blood vessels and blood vessels containing human DPSC-derived endothelial cells (Mean±SEM n=30 and 35 blood vessels, respectively for n=4 grafted animals, ***p=0.0001, one-tailed Mann Whitney test), Scale bar 20 μm.

## DISCUSSION

In this work, we reported for the first time that human DPSCs can be grown using serum-devoid Neurocult proliferation media. In these conditions, both NSCs and DPSCs grew as spheres in suspension. Noteworthy, in DPSCs, this medium also increased the proportion of CD31-positive cells fully viable after one month post-intracraneal graft into nude mice. The vasculature containing human cells expressed VEGF and had increased levels of laminin staining, suggesting that grafted DPSCs favored a process of angiogenesis and neovascularization. These results open a window to use the DPSCs as a safe source of neural crest stem cells to generate endothelial cells which integrate safely into the host brain tissue, approaching new autologous cellular therapies. To our knowledge, this is the first time in which DPSCs disaggregated from dentospheres and grafted into the brain of immunosuppressed mice give rise to CD31-positive endothelial cells that can functionally integrate into brain vasculature, promoting *de novo* generation of new blood vessels within brain tissue. Previous works using myocardial infarction model showed that DPSCs grafts were able to induce heart angiogenesis^43^. Finally, other studies also reported a positive effect of DPSC transplants on the recovery of sensorimotor brain functions after stroke, which was attributed to diverse paracrine mechanisms^44^. However, in all these cases DPSC had been grown in the presence of 10-20 % fetal serum.

Previous authors described the capability of sorted DPSCs grown in the presence of 10% serum to differentiate synergically into osteoblasts and endotheliocytes^18^, mantaining their fate when grafted subcutaneously *in vivo*^18^. However, woven bone chips produced *in vitro* were necessary, contrary to the present results where no bone whatsoever was generated after the *in vivo* DPSC brain graft. Remarkably, it is known that at a 10 - 20 % concentration fetal bovine serum (FBS) can also stimulate osteoblastic differentiation, and this one in turn promotes the generation of neovasculature^45^, although bone production is not desirable in the brain. FBS is added to culture medium to support cell survival and facilitate DPSC expansion. This addition imposes at least two big caveats that make serum use very impractical to apply to DPSC transplants to treat brain lesions: i) the non-desired osteoblastic differentiation of the grafted cells, and ii) the stem cell incorporation and contamination of xenogenic FBS that might cause a dangerous brain inflammatory reaction^23^. In the present work, we employed Neurocult serum-free culture media, which is routinely used for *in vitro* amplification of NSCs and progenitors and do not cause graft rejection.

Very few works in the literature describe the culture of dental stem cells without the use of fetal serum in none of the culture phases: expansion and/or differentiation. Hirata et al. demonstrated for the first time the possibility to maintain stem cells from deciduous or wisdom tooth pulp cells without the need of serum, but without further characterization^46^. Also, Bonnamain et al. were able to culture human pulp stem cells in a serum-free supplemented basal medium, but they reported a 30 % of efficiency to generate non-adherent spheroid-like morphologies^14, 47^. By a protocol using Neurocult proliferation media supplemented with EGF/FGF2, heparin, and B27 without vitamin A, we achieved a near 100 % efficiency in dentosphere generation in DPSC cultures. Previous literature about DPSC intracranial grafts had mainly focused on searching for neuronal fates of the transplanted cells^11, 16^. Our work is the first report describing that non-engineered or non-genetically modified human DPSCs can differentiate and integrate into mouse intracerebral vasculature, promoting neovasculogenesis. However, no sign of NeuN/DCX or GFAP staining was found in the DPSC grafts using Neurocult proliferation media (data not shown).

Nestin expression had been related to *de novo* formation of vasculature^33^, and together with VEGF coexpression in grafted human DPSC-derived cells, it constitutes a solid evidence of neoangiogenesis^48^. Altogether, Nestin/VEGF and/or CD31 human positive cells integrated into the host vasculature with increased laminin staining are the hallmarks that led us to advocate for *de novo* angiogenesis^49, 50^. We could speculate that grafts of human DPSC cultured with Neurocult proliferation media could favor a rejuvenation-like benefitial effect in brain vasculature, because laminin is one of the main extracellular matrix components that progressively decreases up to 50 % during ageing^51^. Interestingly, there is also an increase of vascular TGF-β during ageing, which has been reported to induce quiescence of neurogenic niches^26^. Remarkably again, laminin has been described to be able to downregulate TGF-β, at least in epithelial cells^52^. Thus, the increase of vascular laminin could again constitute a rejuvenation-like stimulus through TGF-β. Furthermore, from a point of view of biological or therapeutic interest, the increase of vascular laminin observed in blood vessels containing DPSC-derived cells could have also beneficial effects in the case of neurodegenerative illnesses such as Alzheimer’s disease. It has been reported that laminin is able to induce depolymerization of amyloid αβ fibrils reducing the toxic effects of amyloid peptides^53, 54^. In cerebro-vascular accident models such as focal cerebral ischemia, it has been reported a degradation of microvascular matrix of collagen, perlecan and laminin^55^. In addition, variations of laminin levels during CNS injury have been associated to an attempt to revascularize and oxygenate the tissue^55, 56^. In other models such as the middle cerebral artery occlusion (MCAO), the extracellular matrix (ECM) and secreted factors such as VEGF are involved in the progressive recovery^55^. *Per se*, the use of DPSCs as a source of cells secreting neuroprotective BNDF also brings an enormous potential for vasculogenesis^57^ and neuroprotective therapies^25, 58^. Interestingly we showed that DPSCs cultured with Neurocult proliferation media maintained Bdnf expression. However, all these discussed applications are beyond the scope of this manuscript, but warrant further investigations.

In this work, we wanted to highlight the possibility to obtain human endothelial cells out of DPSCs that are cultured without genetical modifications or any need of fetal serum at any time. These results are of major interest because no report exists to date which is focused on obtaining endothelial cells starting from DPSCs and using a serum-free protocol. Previous works were focused towards neural differentiation^14, 15, 46^, or endothelial vascular cells using 10 % FBS^18, 59^. Non-human vascular cells had already been succesfully cultured without the need of serum, although the reported protocol was not optimized for human cells^59, 60^. Interestingly, DPSC expansion in Neurocult Proliferation culture media increased the proportion of cells expressing the endothelial markers VEGF and CD31, with respect to neural-differentiated or DMEM/FBS DPSC cultures. DPSC-derived endothelial cells were able to integrate into the murine host brain vasculature increasing levels of laminin, an extracellular matrix component which is progressively lost during aging^51^. In contrast, murine NSC were not able to differentiate towards endothelial cells in the same medium. Some works have reported the need of estradiol for endothelial differentiation of neural progenitors^61^.

Other dental stem cell types such as SHEDs were also reported to be able to differentiate to neural and astroglial lineages^5^ and endothelium^32, 37^. Interestingly, the expression of VEGF by SHEDs was required to induce cellular expression of endothelial differentiation markers such as vascular endothelial growth factor receptor-2 (VEGFR2), CD31 (PECAM-1) and VE-Cadherin^32^. Similar results have been published with stem cells from the apical papilla (SCAPs) or DPSC previously expanded with α-MEM^18, 19^. However, once again all the aforementioned studies relied on the presence of 10% FBS in the cell culture medium, and/or the use of hydroxyapatite/tricalcium phosphate 3D scaffolds to show the successful formation of vascularized tissue^32^. Our results demonstrate that a serum-free medium designed to culture neural stem cells is permissive to grow endothelial cells derived from human DPSC cultures.

## METHODS

### CELL CULTURE AND CELL PROLIFERATION

Human third molars were obtained from healthy donors of between 15 and 45 years of age. Tooth samples were obtained by donation after informed consent, in compliance with the 14/2007 Spanish directive for Biomedical research, and the protocol was approved by the CEISH committee of UPV/EHU. DPSC isolation and culture were carried out as previously reported ^4^. Briefly, DPSCs were isolated by mechanical fracture and enzymatic digestion of the pulp tissue for 1 h at 37 °C with 3 mg/ml collagenase (17018-029, Thermo Fisher Scientific, Waltham, MA USA), and 4 mg/ml dispase (17105-041, Thermo Fisher Scientific, Waltham, MA USA). After centrifugation at 1500 rpm for 5 minutes, cells were resuspended and underwent mechanical dissociation by 18 G needles (304622, BD Microlance 3). Then DPSCs were cultured in parallel with different types of culture media: (i): DMEM (Lonza 12-733, Basel, Switzerland) supplemented with 10 % of inactivated FBS (SV30160.03, Hyclone, GE Healthcare Life Sciences, Logan. UT, USA), 2 mM L-Glutamine (G7513, Sigma, St. Louis, MO) and 100 U/mL penicillin + 150 μg/mL streptomycin antibiotics (15140-122, Gibco). (ii): Human Neurocult medium composed of Human Neurocult NS-A basal medium (cat# 05750, Stem Cell Technologies, Vancouver, Canada) with Neurocult proliferation supplement (cat# 05753, Stem Cell Technologies, Vancouver, Canada) or Neurocult differentiation supplement (cat# 05752, Stem Cell Technologies, Vancouver, Canada) both at 9:1 ratio, and supplemented with Heparin solution 2 µg/ml (cat# 07980, Stem Cell Technologies, Vancouver, Canada), EGF 20 ng/ml and FGFb 10 ng/ml (Peprotech, London, UK) as previously described^26^ in presence of antibiotics penicillin 100 U/ml and streptomycin 150 µg/ml (15140-122, Gibco). For NSC cultures isolated from Nestin-GFP mice, dissected hippocampi were removed with ice-cooled PBS-sucrose and processed as SVZ as previously described^26^. Cells were maintained at standard conditions in a humidified 37 Cº incubator containing 5% CO_2_. Neurosphere cultures were then passaged every 7 days by enzymatic disaggregation with Accutase (Sigma, St. Louis, MO). We cultured DPSCs cells for 1 month and a maximum of 4 total passages in order to avoid cell aging issues.

The population doubling (PD) rate was determined by initial cell culture. Cells, neurospheres and dentospheres were disaggregated, counted and passaged at day 7. At each passage cells were re-plated at the initial density and cultures were performed until passage 4. The population doubling rate was calculated using the following formula:

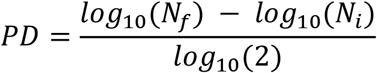

*N*_f_ is the harvested cell number and *N*_i_ is the initial plated cell number. Cumulative population doublings (CPD) index for each passage was obtained by adding the PD of each passage to the PD of the previous passages as previously described ^62^. All cells were seeded at the density of 6 × 10^3^ cells/cm^2^ and cultured for 1 week. Cell counting was performed after cell detachment or dissociation using an automated TC20 from Bio-Rad cell counter. The total number of cells estimation was calculated on three experimental samples for each type of culture.

### FLOW CYTOMETRY

Half-million DPSC grown in either DMEM 10% FBS or Neurocult proliferation media were detached, disaggregated and then incubated with PBS 0.15% BSA solution with 0.5 μg of CD31-FITC or IgG2a κ Isotype control for 1h at 4°C. After a wash with PBS 0.15% BSA cells were resuspended in 300 μL of PBS 0.15% BSA and analyzed using a FACS Beckman Coulter Gallios (Beckman Coulter Life Sciences, Indianapolis, United States). The data were analyzed using Flowing Software 2.5 (University of Turku, Finland).

### ANIMALS AND CELL GRAFT

Consanguine c57bl6 litters from Nestin GFP mice and Athymic Swiss^nu/nu^ were used as hosts for murine and human *in vivo* graft purposes. Nestin-GFP neurospheres or DPSCs dentospheres in the active growth phase were disaggregated, washed and collected in Neurocult serum-free media. Two microliters containing 100,000 cells were injected (0.5 µL/min) unilaterally at the following coordinates (from bregma): AP = −1.9, L = −1.2, and DV = −2 and −2.1. The cell transplantations were performed using a small animal stereotaxic apparatus (Kopf model 900) with a 10 µl Hamilton syringe and a 33 G needle (Hamilton, Bonaduz, Switzerland). Pre-operatory and post-operatory animal care were carried out as previously described^42^. Animals were provided with food and water ad libitum and housed in a colony isolator maintained at a constant temperature of 19–22 °C and humidity (40–50%) on a 12:12 h light/dark cycle. The animal experiments were performed in compliance with the European Communities Council Directive of November 24, 1986 (86/609/EEC) and were approved by the competent authority (Administración Foral de Bizkaia).

### IMMUNOSTAINING OF BRAIN SECTIONS AND CELL CULTURE

Animals were deeply anesthetized with Avertin 2.5 % and transcardially perfused with a 4 % paraformaldehyde solution in 0.1 M sodium phosphate, pH 7.2 and processed as previously described^26^. In order to detect grafted genetically unmodified human DPSCs on mice brain, specific antibodies targeted to human Nestin (MAB1259, 1:200 R&D systems)^42^, and human CD31 (BBA7, 1:200 R&D systems) were used. Immunostaining of brain vasculature was developed using CD31 (550247, 1:300 BD Pharmingen), laminin (L9393, 1:200 Sigma, St. Louis, MO), and VEGF (ABS82-AF647, 1:200 Sigma, St. Louis, MO) antibodies, and immunostaining of macrophages/microglia with CD68 antibody (MCA-1957, 1:400 BioRad). For cell culture both NSCs and DPSCs, either in the form of dissociated cells or small spheres, were seeded into laminin-treated coverslips (L2020, Sigma, St. Louis, MO) as previously described^63^. After three days or one week, they were fixed by incubation with 4 % PFA for 10 minutes at room temperature and permeabilized by incubation in 0.1 % Triton X-100. They were then incubated overnight at 4 ºC with primary antibodies at the following dilutions: Glial Fibrillary Acidic Protein (GFAP) (MAB3402, 1:400; Millipore, Lake Placid, NY), Nestin (NES, 1:200 Aves Labs), S100ß (Z0311, 1:1000, Dako, Glostrup, Denmark), NeuN (EPR12763, 1:200; Abcam, Cambridge, UK), Doublecortin (DCX) (sc-8066, 1:200; Santa Cruz, Dallas, TX, USA), CD31 (550247, 1:300 BD Pharmingen, San Jose, CA, USA), VEGF (ABS82-AF647, 1:200 Sigma, St. Louis, MO), laminin (L9393, 1:200 Sigma, St. Louis, MO). For both tissue sections and cell culture, secondary antibodies conjugated to Alexa 488, 568 and 647 Donkey anti-mouse, anti-rabbit or anti-goat were incubated for 2 h and 30 min respectively at room temperature. Preparations were counterstained with DAPI and images were captured using a Leica SP8 confocal microscope at 40X magnification.

### CONVENTIONAL PCR AND QUANTITATIVE REAL-TIME PCR (QPCR)

RNA extraction from cell pellets, reverse transcription and QPCR were performed as previously described^64^. The molecular weights of the amplification products were checked by electrophoresis in a 2 % agarose gel. All reactions were performed in triplicate and the relative expression of each gene was calculated using the standard 2^−ΔΔ^Ct method^65^. Primer pairs used were obtained through the Primer-Blast method (Primer Bank) and they are listed in Table 1.

### WESTERN BLOT

DPSCs either grown using NeuroCult proliferation medium or DMEM + 10% FBS and Liver Sinusoidal Endothelial Cells (LSEC) were counted and resuspended in a ratio of 20,000 cells /μl of RIPA lysis buffer (R0278, Sigma, St. Louis, MO) supplemented with protease (11873580001; Roche) and phosphatase inhibitors (78420; Thermo Scientific) to ensure the same cellular concentration for the different type of cells and culture media. From this, thirty micrograms of protein was diluted in RIPA buffer supplemented with LDS Sample Buffer (NP0007; Invitrogen by Life technologies). Electroblot was performed as previously described^42^. VEGFR, phosphorylated-ERK and total ERK antibodies were used to detect the VEGFR signaling pathway (1:1000, #9698S, #4370 and #4695 respectively, Cell Signaling Technologies). Beta-actin (1:1000, A5441, Sigma St. Louis, MO), Stat3 (1:1000, #9132, Cell Signaling Technologies) and Ponceau staining (P7170-1L, Sigma St. Louis, MO) were used as loading control to detect protein in the charged lanes.

### STATISTICAL ANALYSIS

Comparisons between multiple groups were made using Kruskal Wallis followed by Dunn’s post hoc test. Comparisons between only two groups were made using U-Mann Whitney test or Student’s t test. p<0.05 was considered as statistically significant. Results were presented as mean ± SD or SEM. The number of independent experiments is shown in the respective section.

### STUDY APPROVAL

The animal experiments were performed in compliance with the European Communities Council Directive of November 24, 1986 (86/609/EEC) and were approved by the competent authority (Administración Foral de Bizkaia).

## AUTHOR CONTRIBUTIONS

J.R.P., F.U., J.M.E. and G.I.: conception and design; J.R.P., G.I., O.P.A., F.U. and J.L.: provision of study materials and manuscript writing; J.R.P., J.M.E., F.U. and G.I.: financial support; J.L., O.P.A. and J.R.P.: collection and/or assembly of data, J.L., G.I. and J.R.P., data analysis and interpretation; J.L., O.P.A., J.M.E., F.U., G.I. and J.R.P., manuscript discussion and final approval of manuscript.

## ACKNOWLEDGMENTS

Authors would like to thanks to all the staff of the Leioa animal facility of UPV/EHU, Escobar L. and Díez A. for technical support of microscopy and for flow cytometry services in the Achucarro and SGIKER of UPV/EHU. Liver sinusoidal endothelial cells (LSEC) were a generous gift from Prof. Iker Badiola. This work has been funded by « Ramón y Cajal » program RYC-2013-13450 (JRP) and RYC 2012-11137 (JME); Spanish Ministry of Economy and Competitivity SAF2015-70866-R; UPV/EHU (GIU16/66, UFI 11/44) and Basque Government (GV/EJ; IT831-13). JL and OPA obtained a Ph.D. fellowship from the University of the Basque Country (UPV/EHU).

